# Accumulation of a biparentally-inherited Neptune transposable element in natural Killifish hybrids (*Fundulus diaphanus* × *F. heteroclitus*)

**DOI:** 10.1101/2025.05.22.655539

**Authors:** Alexandre-James Roussel, Alexander Suh, Francisco J. Ruiz-Ruano, Anne-Marie Dion-Côté

## Abstract

Transposable elements (TEs) are abundant selfish genetic elements that can mobilize in their host genome, causing DNA damage, mutations and chromosome rearrangements. TE silencing is thus critical, and is initiated by maternally loaded piRNAs, leading to their repression. Consistently, paternally inherited TEs are derepressed in the progeny of Drosophila crosses involving a naive female. TEs have also been found to be derepressed in interspecific crosses, which is proposed to result from suboptimal interactions of piRNA pathway proteins. *Fundulus heteroclitus* and *F. diaphanus* hybridize in nature and produce viable and fertile offspring that sometimes reproduce asexually. We characterized the repetitive DNA content of these species and their asexually reproducing hybrids. TE load was slightly higher than expected in hybrids and associated with younger repeats. Two bi-parentally inherited active Neptune subfamilies showed a remarkable ∼3-4-fold accumulation in hybrids. These results are consistent with suboptimal piRNA pathway function, leading to active TE accumulation.

## Introduction

Hybridization between divergent populations or species may trigger genome instability in hybrids, a phenomenon long proposed to be associated with transposable element (TE) derepression (i.e., the genomic shock hypothesis, McClintock 1984). TEs are selfish repetitive sequences that can mobilize within their host genome and are classified in two major groups. Class I elements (retrotransposons) require an RNA intermediate which is retro-transcribed into complementary DNA before re-inserting in the genome, increasing copy number. Retrotransposons are further classified into subclasses, depending on the presence or absence of long terminal repeats (LTRs and non-LTRs) (Wicker et al. 2007). *Penelope*-like elements (PLEs) form a peculiar type of non-LTR retrotransposons due to their unique organization and mode of mobilization, and the presence of a unique GIY-YIG endonuclease domain (Evgen’ev and Arkhipova 2005; Frangieh et al. 2024). Class II elements (DNA transposons) do not transpose by reverse transcription but rather via DNA excision and insertion (Wicker et al. 2007).

TE mobilization can be highly deleterious, disrupting genes or regulatory elements, and inducing DNA damage or chromosome rearrangements (Bourque et al. 2018). To counteract these threats, silencing mechanisms have evolved in eukaryotes to repress TEs (Cosby et al. 2019). In animals, a key component of this defense system involves Piwi-interacting RNAs (piRNA) and the associated piRNA pathway proteins, which together mediate TE repression (Le Thomas et al. 2013). In Drosophila, this silencing is initiated by maternally transmitted piRNAs, safeguarding the progeny’s genome against TEs (Malone et al. 2009). Importantly, piRNA pathway proteins themselves evolve rapidly, a pattern that is widely interpreted as a response to the fast-paced sequence evolution of selfish TEs. This dynamic reflects an ongoing evolutionary arms race, where host silencing machinery must continually adapt to recognize and suppress newly emerging or mutating TEs evading repression (Blumenstiel et al. 2016; Cosby et al. 2019; Parhad and Theurkauf 2019).

Observations across various systems have led to two non-mutually exclusive models that may account for TE mobilization in hybrids. The first model involves paternal-only TE derepression, where active TEs transmitted by the paternal line but absent from the maternal genome escape silencing and mobilize in hybrids. This phenomenon is thought to result from the lack of maternally-transmitted piRNAs that match and silence paternally inherited TEs. This phenomenon is well-documented in Drosophila dysgenic crosses but has not been thoroughly descrived in other systems, to the best of our knowledge (Bucheton et al. 1976; Kidwell et al. 1977; Evgen’ev et al. 1997, Marie et al. 2017, Ryazansky et al. 2017). In contrast, the second model involves bi-parental TE derepression, where TEs inherited from both parents are activated. It has been proposed that bi-parental TE rerepression results from the accumulation of genetic incompatibilities in piRNA pathway proteins over evolutionary time, which impairs TE silencing mechanisms in interspecific hybrids (Kelleher et al. 2012). Such bi-parental TE derepression has been reported in Drosophila interspecific hybrids (Labrador et al. 1999; Kelleher et al. 2012; Romero-Soriano et al. 2017), as well as in some mammal, plant and fish hybrids (O’Neill et al. 1998; Ungerer et al. 2006; Dion-Côté et al. 2014). While TE derepression has been observed in various aforementioned hybrid systems, its downstream genomic impact remains unclear. In some cases, studies have focused solely on transcriptional derepression, leaving open the question of whether increased TE expression leads to greater TE accumulation or transposition. Moreover, many studies rely on later-generation hybrids, making it difficult to determine whether derepression occurred in the initial F_1_ or F_2_ generations, or emerged later. In other cases, the observed genomic instability may be confounded by chromosomal rearrangements or the effects of meiotic recombination, which cannot be excluded as alternative or contributing mechanisms.

*Fundulus heteroclitus* and *F. diaphanus* are killifish species that diverged between 15 and 25 million years ago (Ghedotti and Davis 2017). While *F. diaphanus* is primarily found in freshwater, *F. heteroclitus* preferentially occurs in saltwater (Scott and Crossman 1973). Both species are euryhaline and co-occur in brackish water, where sexually and asexually reproducing hybrids have been reported (Fritz and Garside 1974; Dawley 1992; Hernández Chávez and Turgeon 2007; Mérette et al. 2009). An opening formed ∼70 years ago (∼25-40 generations) in Porters Lake, NS, allowing sea water to enter (Fritz and Garside 1974). Therefore, secondary contact between *F. heteroclitus* and *F. diaphanus* in Porters Lake is very recent, from ∼70 years to a conservative maximum of 13 000 years, as the area was covered by Wisconsinian glaciers until then (Shaw et al. 2006; Dalziel et al. 2020). There, previous work has revealed the presence of asexually reproducing “frozen F1” hybrids that reproduce by gynogenesis, carrying a full, unrecombined chromosome set from each parental species due to disrupted meiotic recombination between homologs (Hernández Chávez and Turgeon 2007; Dalziel et al. 2020, Dion-Côté et al., *unpublished observations*). Several clonal lineages co-exist, but one lineage is predominant (Hernández Chávez and Turgeon 2007; Dalziel et al. 2020). This system provides a rare opportunity to investigate maternally- and paternally-transmitted TE accumulation in “frozen F1” hybrids, controlling for some of the confounding effects of meiotic recombination and large-scale chromosome rearrangements. The relatively young age of the *Fundulus* clones also allows to capture TE accumulation in these hybrids, if any, before their decay, as predicted under long-term asexual reproduction (Hickey 1982; Arkhipova and Meselson 2005; Dolgin and Charlesworth 2006).

## Material and Methods

### Fish collection and sampling

Experimental procedures were approved by the Université de Moncton’s animal care and use committee (19-02). Fishes were collected from Porters Lake, Nova Scotia (44°43′40″N, 63°18′00″W) during Summer 2020 using beach seine nets under a Department of Oceans and Fisheries licence (355651). Because asexually reproducing hybrids are all females, only females were kept (Table S1). Fish were housed in brackish water (12 ppt) at Université de Moncton until they were sacrificed. Fin clips were preserved in 95% ethanol.

### DNA extraction, genotyping and sequencing

DNA from three *F. heteroclitus*, four *F. diaphanus* and seven hybrid individuals was extracted from fin clips using the E.Z.N.A tissue DNA extraction kit (Omega Bio-tek). To determine maternal ancestry, a 441bp fragment from the mtDNA D-Loop was amplified by PCR and digested with HphI (New England BioLabs), as previously described (Dalziel et al. 2020). In addition, five microsatellite loci were genotyped for each individual [FhATG-B103, FhCA-1 and FhCA-21: (Adams et al. 2005), Fhe113 and Fhe57: (Hernández Chávez and Turgeon 2007), Table S2]. Allelic sizes were determined by capillary electrophoresis on a 3730xl DNA Analyzer (Applied Biosystems) at The Center for Applied Genomics, Toronto. Results were analyzed using the Microsatellite Analysis application (ThermoFisher Connect Cloud).

PCR-free DNA libraries (Lucigen) were prepared and sequenced on an Illumina HiSeq Instrument with 150 bp paired-end reads at the Genome Quebec Innovation Center (Table S3). Quality control was performed using FastQC (V0.11.9). Reads were trimmed with Trimmomatic (v0.38) using the following parameters: ILLUMINACLIP 2:30:7:4, HEADCROP: 10 and MINLEN: 100.

### Repeat characterization

We characterized the most abundant repeats in hybrid individuals in a single run of RepeatExplorer2 (Novák et al. 2013). We randomly selected and concatenated 71,429 trimmed read pairs from each of the seven hybrid individuals for a total of 500,000 read pairs. These reads were grouped by RepeatExplorer2 into clusters. We further classified each cluster into the most specific group possible, based on a hierarchical decision tree (Fig. S1). For TEs, a cluster is equivalent to a complete or partial subfamily. “Unclassified repeats” are clusters that were not annotated as a satellites, mobile elements or rDNA, whereas “Unknown” clusters were not classified as organelles or contamination. The final repeat database was composed of 14,011 contigs (Supplementary File 1).

We used the “id_rmasker.py” script to add the annotation to the sequence IDs in RepeatMasker’s format (Smit et al. 2013-2015) and the “replace_patterns.py” script to transform the RepeatExplorer2 annotation to be cluster-specific (https://github.com/fjruizruano/ngs-protocols). We used the resulting repeat database to estimate the proportion of the genome occupied by each cluster. To do this, we selected 5 million trimmed read pairs per sequenced library using seqtk v1.3 (https://github.com/lh3/seqtk). We then quantified the abundance of each repeat within the database with this subset of reads using RepeatMasker v4.1.3 (Supplementary Files 2–3). RepeatMasker was run using the repeat_masker_run_big.py script from the satMiner toolkit (https://github.com/fjruizruano/satminer, last access on 11/07/2022) (Ruiz-Ruano et al. 2016). This script runs RepeatMasker on large query files by splitting them up into subsets. It also runs the calcDivergenceFromAlign.pl script from RepeatMasker, which counts the total number of nucleotides occupied by each group of repeats along divergence intervals (Kimura substitution levels) of 1% needed to generate repeat landscapes. Genome proportion occupied by each cluster was obtained by normalizing the number of nucleotides occupied by each cluster to the total number of nucleotides for its subset of reads. We then generated average repeat landscapes for each group (*F. heteroclitus, F. diaphanus*, and hybrids) and subtractive repeat landscapes as the average values of a group minus the average values of the other group. A principal component analysis was run on the abundance for each repeat category using the function PCA() from the FactoMineR package (Lê et al. 2008) in R. We calculated the average genome proportion of the parental species and the hybrids for each cluster. We then detected important deviations from the observed genome proportion occupied by each repeat cluster in hybrids compared with the expected proportion as the average between the genome proportion of both parental species. This analysis revealed repeat.CL27 as being ∼4X more abundant than expected.

### Haplotype calling

We tested whether some species-specific subfamilies of repeat.CL27 preferentially accumulated in hybrids using the sequence of ORF2 (reverse transcriptase, see *Copy number validation by qPCR*) from the repeat.CL27 consensus sequence as a reference. We mapped the trimmed paired reads from the 14 individuals using bwa-mem2 v.2.2.1 with relaxed conditions (“-SP -k 14 -L 0,0 -U 0 -B 1”). We processed the resulting BAM files with samtools v1.14 (Danecek et al. 2021) as follows. First, we sorted and kept reads mappings with MAPQ ≥ 40 and merged the mapped reads for each parental species separately. We then added the read group for each BAM file with Picard v3.1.1 (Broad Institute). We performed haplotype calling with freebayes v1.3.2 using a minimum length of 7 (--haplotype-length 7) (Garrison and Marth 2012). We extracted haplotypes from the resulting VCF file with BCFtools query v.1.10.2 (Danecek et al. 2021) applying the option (-f ‘%CHROM\t%POS\t%REF\t%ALT\t[%SAMPLE:%GT\t]\n’). Then, we extracted private haplotypes from each parental species and selected haplotypes longer than 15 nt. Finally, we extracted the mapped reads from the 14 sequencing libraries (parental species and hybrids) in FASTQ format with samtools fastq and counted the number of occurrences of each haplotype with grep (Supplementary File 4).

### Copy number validation by qPCR

We identified open reading frames (ORF) in the consensus sequence of repeat.CL27 (CL27Contig27_13_sc_0.609135_l_5617) using ORFfinder (https://www.ncbi.nlm.nih.gov/orffinder/), which identified two long non-overlapping ORFs (ORF1 and ORF2, 420 and 2004 nt, respectively). We then estimated the number of copies in 18 individuals (six individuals for each group) by qPCR. We designed primers targeting the ORF2 from repeat.CL27 using Primer-BLAST (F: 5’-AGGAACCTGAAAAGTGGAGCA-3’, R: 5’-TGGTGCCGAATTGAGAGAGG3’) and the citrate synthase gene from the *F. heteroclitus* reference genome (NC_046361, F: 5’-TGCACCCCATGTCTCAGTTC-3’, R: 5’-TTGGACTTGTGCACTCCCTC-3).

Efficiency curves were generated using 0.0032 - 10 ng (5-fold serial dilution) of gDNA from a single *F. heteroclitus* female individual. Reactions (7 μl) contained 0.25 μM of each primer and 1X Luna Universal qPCR master mix (New England BioLabs). Amplification began with an initial denaturation at 95°C for 1 min, followed by denaturation at 95°C for 15 sec and combined annealing and amplification at 60°C for 30 sec, for a total of 40 cycles. Amplification efficiency was acceptable for both genes (97.3%, R^2^ = 0.994 for citrate synthase, 100.65%, R^2^ = 0.994 for ORF2 of repeat.CL27) and melting curves revealed a single peak per reaction. Both reactions were run on the same plate for each sample.

### Cloning and sequencing of repeat.CL27 ORF2

We manually designed primers to amplify the full-length reverse transcriptase encoded by ORF2 from the consensus sequence of repeat.CL27 (F: 5’-ATGAGGTATGACCTGCAAAAATACC-3’, R: 5’-GATCCAAATCCTTTCAGCCCTTC-3’). The reactions (25 µl) contained 0.6 µM of each primer, 2 mM MgCl_2_, 2 mM dNTPs, 1.25 U GoTaq and 45 ng of DNA. The amplification protocol included an initial denaturation step at 95°C for 2 min, followed by 32 cycles of denaturation at 95°C for 1 min, annealing at 60°C for 1 min and amplification at 72°C for 1 min, with a final elongation at 72°C for 5 min. Amplicons were gel-purified using the Monarch DNA gel extraction kit (New England BioLabs). Purified amplicons were cloned using the TOPO TA cloning kit for sequencing (Invitrogen) following the manufacturer’s recommendations and transformed in DH5a competent cells (Fisher Scientific). Sequencing of the plasmids was performed by Plasmidsaurus, and vector sequence was removed using A Plasmid Editor (Davis and Jorgensen 2022) (Supplementary file 5).

## Results and Discussion

*F. diaphanus* had a slightly higher repetitive DNA load than *F. heteroclitus* (30.39% vs 29.30%, p < 0.001, Type III ANOVA, Table S4), consistent with its larger genome (1.48 Gb for *F. diaphanus* vs 1.33 Gb for *F. heteroclitus*, Dawley 1992). For *F. heteroclitus*, the proportion of the genome occupied by TEs was lower than previously reported using an assembly-based approach, which is more sensitive to low copy number repeats in the assembled genome (32.66%, Reinar et al. 2023). In fishes, TEs are highly diversified, with non-LTR retrotransposons and DNA transposons typically being the most abundant, with many families that remain active (Volff 2005; Shao et al. 2019). As expected, non-LTR retrotransposons were the most abundant TEs in both species. On the contrary, we identified very few DNA transposons (<1% of the genome for Maverick elements and no hATs), but this would be expected if many of them were non-autonomous, i.e., lacking any ORFs (Peona et al. 2024). We compared the repetitive DNA content of *F. diaphanus* and *F. heteroclitus* by generating repeat landscapes which revealed higher abundances among the lower Kimura substitution level bins for both species, consistent with a recent accumulation of repeats (Fig. 1A & B). A subtractive repeat landscape, calculated as *F. diaphanus* minus *F. heteroclitus*, revealed that the main differences between both species were found among the less divergent repeats, i.e., those that expanded more recently (Fig. 1C). This analysis also revealed an expansion of many TE groups as well as unknown/unclassified repeats in *F. diaphanus* (or a contraction in *F. heteroclitus*) whereas young satellite DNA appeared to have recently expanded in *F. heteroclitus* (or contracted in *F. diaphanus*).

**Figure 1.**
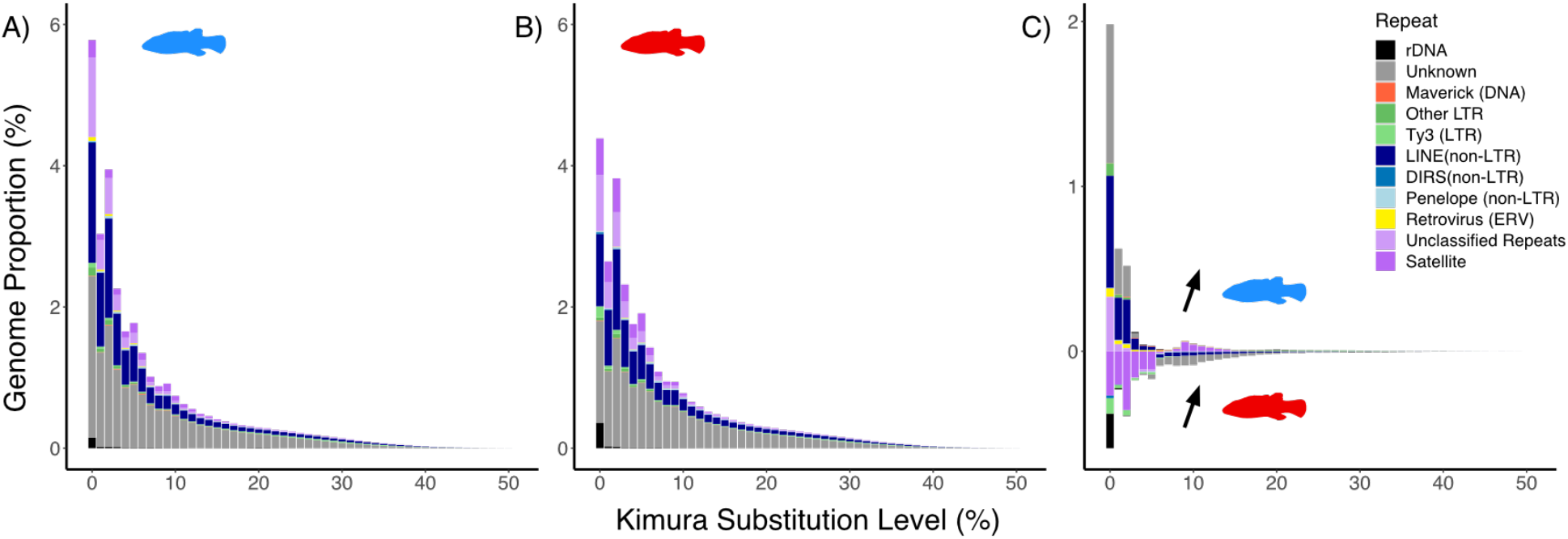
Repeat content in *Fundulus* spp. We calculated the genome proportion occupied by each repeat group as a function of the Kimura substitution level (CpG corrected) of a given repeat cluster to its consensus sequence. A) Repeat landscape of *F. diaphanus* (average content across four individuals). B) Repeat landscape of *F. heteroclitus* (average content across three individuals). C) Subtractive repeat landscape generated by subtracting *F. heteroclitus* (B) from *F. diaphanus* (A) average abundance values. Annotation based on Fig. S1. Silhouette from PhyloPic.

To visualize how repeat content in hybrids differs from the parental species, we performed a principal component analysis (PCA) on the genome proportion occupied by each repeat group. Principal components (PC) 1 (66.1% of the variance) clearly separated parental species, which was expected given their contrasted repeat landscapes (Fig. 2A). Along PC1, hybrids were also intermediate between parental species, although slightly closer to *F. diaphanus*. This was expected as *F. diaphanus* has a larger genome. PC2 (25.4% of the variance) clearly separates hybrids from parental species, with all repeat groups being positively correlated to this PC, except for rDNA (more abundant in *F. heteroclitus*) and retroviruses (more abundant in *F. diaphanus*). This suggests that most repeat groups accumulated in hybrid genomes. We thus generated a subtractive repeat landscape to compare the observed hybrid repeat content with the “expected” repeat content (i.e., the sum of half the average abundance of each parental species for each repeat group, Fig. 2B). This revealed that hybrids had a higher repetitive DNA content than expected for every repeat group, except rDNA (Fig. 2C). Yet, the genome proportion occupied by repeats in hybrids was only 0.665% higher than in *F. diaphanus* and 1.69% in *F. heteroclitus* (Table S4). These results suggest that most repetitive elements have accumulated in *Fundulus* hybrids, consistent with suboptimal piRNA pathway protein interaction for TEs, although very moderately. While we cannot entirely exclude technical biases, such as underestimating hybrid genome size or overestimating repeat abundance through parental allele separation, this pattern would not be restricted to the youngest repeats (i.e., lowest Kimura substitution bins) as we observe. Another caveat is that the individuals from the parental species that we sequenced may not fully capture the variation present in the parental individuals that produced the sequenced clones.

**Figure 2.**
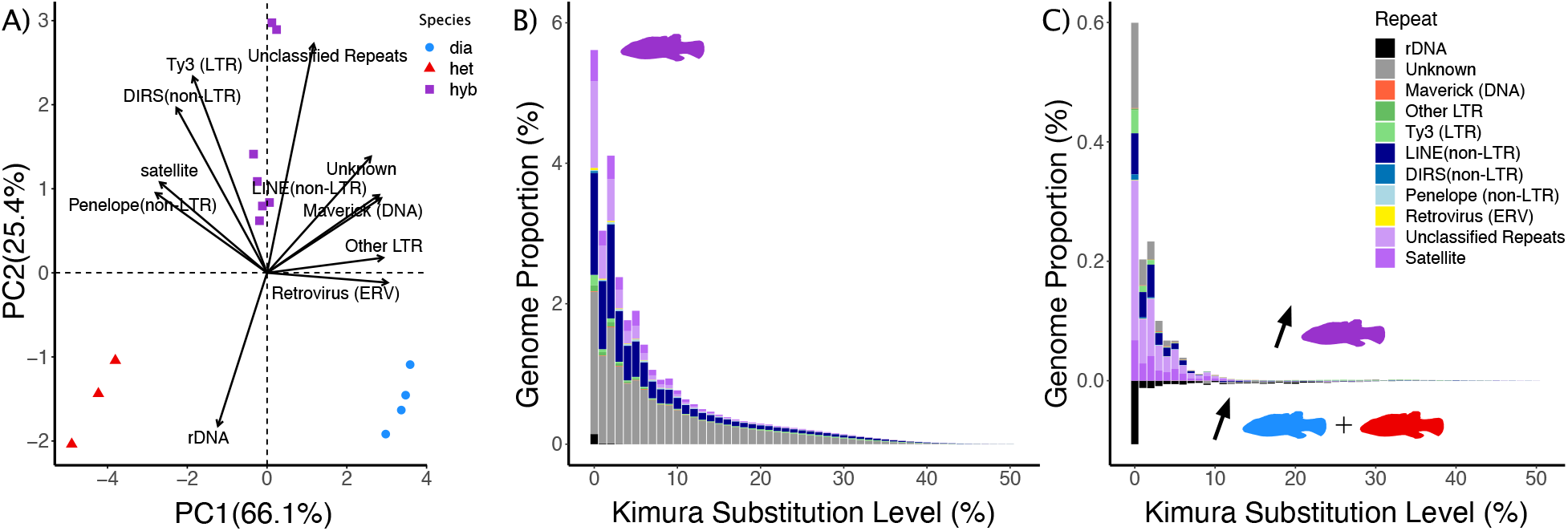
Repeat content in *Fundulus* hybrids. A) PCA of the repeat group content on *Fundulus* fish. B) Repeat landscape of *Fundulus* hybrids. C) Subtractive repeat landscape generated by subtracting expected abundance values (half the average abundance in *F. diaphanus* plus half the average abundance in *F. heteroclitus*) from the abundance observed in hybrids. Annotation based on Fig S1.

Therefore, we examined whether some repeats expanded more extensively in hybrids, compared to our expectation (i.e., the sum of half the average abundance of each parental species for each repeat group, Fig. 3A). We extracted the only three repeat clusters that varied by more than 1 Mb between the observed and expected hybrid repeat content (Fig. 3A). The first one, annotated as a Ty3 element (LTR retrotransposon), was ∼1.15 times more abundant in hybrids than expected, indicating a moderate increase. Although satellite.CL1 was more abundant in hybrids than expected (observed: 1.37 × 10^7^ ± 1.13 × 10^6^ bp, expected: 1.17 × 10^7^ bp), it fell within the *F. heteroclitus* range (1.35 × 10^7^ ± 1.15 × 10^6^ bp). A third candidate, repeat.CL27 (“unclassified”, Fig. S1), stood out, being ∼4 times more abundant than expected in hybrids (Fig. 3B, p < 0.001, Type III ANOVA,). This result was further validated by qPCR with different individuals, to exclude methodological biases, which revealed a 2.92 average increased abundance in hybrids compared to parental species (Fig. S2). Based on CENSOR (Kohany et al. 2006), this repeat presents similarity (positions 5128 to 2560, minus strand) with a Neptune retroelement previously identified in the Threespine Stickleback (*Gasterosteus aculeatus*) (subfamily Neptune-2_GA, positions 5128-2460, 66.03% identity). Elements from the Neptune superfamily are poorly characterized *Penelope*-like elements defined by the presence of a reverse transcriptase (RT) and a GIY-YIG endonuclease domain (Evgen’ev and Arkhipova 2005) that have now been detected across algae, plants, fungi, protists and animals (Craig et al. 2021). We identified two long ORFs in repeat.CL27, from position 73-492 (419 nt, ORF1) and from position 666-2669 (2003 nt, ORF2). We did not detect sequence similarity for ORF1 in any of the queried databases (Repbase, GenBank, gtRNAdb, CDD), but ORF2 contains the intact RT (AA 590-558) and GIY-YIG (AA 299-392) domains based on the Conserved Domains Database (CDD). A phylogeny with other PLEs revealed that ORF2 groups with other Neptune subfamilies (Fig. 3C). We cloned and sequenced full ORF2 from various individuals, which revealed the existence of two subfamilies of this Neptune element that are specific to each parental species (Fig. 3D). Both subfamilies were found in hybrids, which prompted us to examine the accumulation of short, species-specific haplotypes in repeat.CL27 from the sequencing reads (Supplementary File 4). We detected 8 regions > 15 nt, resulting in 27 haplotypes, 12 of which were specific to *F. diaphanus* and 7 to *F. heteroclitus* (i.e., short haplotypes that are diagnostic of a subfamily that is restricted to either parental species). Among *F. diaphanus*-specific haplotypes, five out of seven were more abundant in hybrids than expected (observed abundance/expected abundance median = 6.79). Likewise, five out of 12 *F. heteroclitus*-specific haplotypes showed a higher abundance in hybrids than expected (observed abundance/expected abundance median = 7.12). Thus, some species-specific haplotypes are present in excess in these asexually reproducing “frozen F1” hybrids, whereas others are under-represented. Species-specific haplotypes may be under-represented in hybrid genomes because they were absent in the original parents, emerged in parental species after hybrid formation, or were lost in hybrids through mutations or deletions. Altogether, these results suggest that species-specific variants of this Neptune subfamily from each parental species are accumulating in *Fundulus* hybrids. We propose that this accumulation may be restricted to currently active variants.

**Figure 3.**
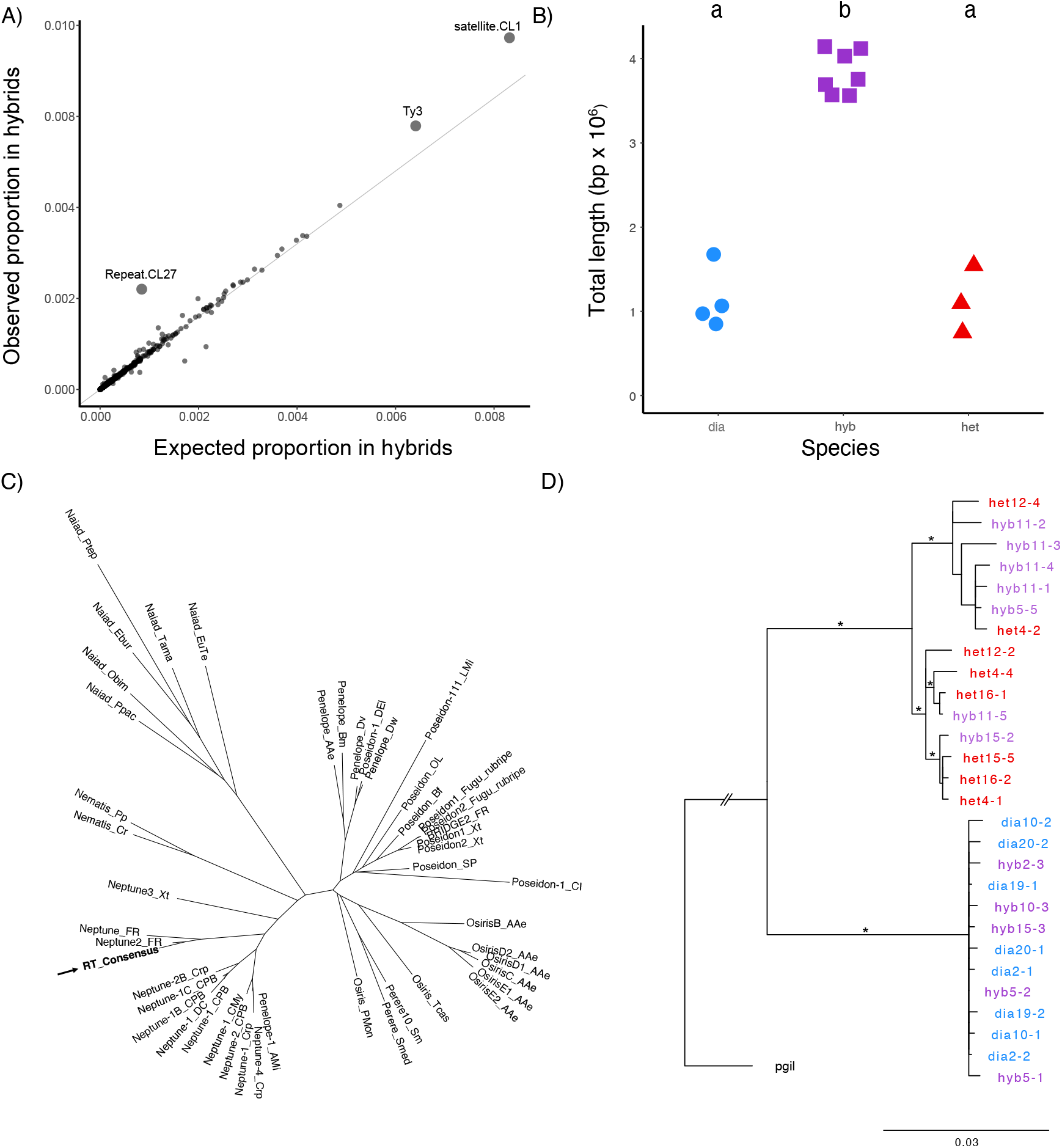
A Neptune transposable element accumulates in *Fundulus* hybrids. A) Observed proportion of the genome occupied by each repeat cluster in hybrids as a function of the expected copy number. Repeat.CL27 is highlighted. The line depicts a 1:1 ratio. B) Total genomic length (bp) occupied by repeat.CL27 based on sequencing (p < 0.001, Type III ANOVA). C) Phylogeny of PLEs RT and repeat.CL27 ORF2 (RT consensus). PLEs sequences were graciously provided by Tomás Carrasco-Valenzuela (Carrasco-Valenzuela et al. 2023). D) Phylogeny of cloned full-length RT from 14 individuals. Outgroup sequence from *Poecilia gillii* (NCBI accession number CAEGAJ010041810.1: 443 - 2446) (van Kruistum et al. 2021) was obtained by doing a BLASTN search of the consensus sequence against the WGS database restricted to the *Poecilia* genus (taxid:8080). Sequences were aligned using MAFFT-LINSI v7.505 (Katoh and Standley 2013) (Supplementary file 6). We manually inspected the resulting alignment for the presence of PCR chimeras and removed them. We built a phylogeny using IQ-TREE v2.4.0 with default parameters and 1000 bootstraps (Nguyen et al. 2015). Trees were edited using Figtree v1.4.4 (https://github.com/rambaut/figtree). Branches with >0.85 bootstrap support are indicated with an asterisk.

## Conclusion

While the moderate accumulation of many repetitive sequences we observed in hybrids is coherent with suboptimal piRNA pathway protein interactions (Kelleher et al. 2012), technical biases cannot be entirely excluded. We identified two Neptune subfamilies, each of which are specific to either *F. diaphanus* or *F. heteroclitus*, that accumulated dramatically in naturally occurring “frozen F1” hybrids. This pronounced accumulation of two TE subfamilies, each inherited from either parental species, differs from previously described patterns: in *Drosophila* dysgenic crosses, where only paternally inherited TEs are derepressed in the progeny of naïve females, and in other interspecific hybrids, where multiple TE families tend to accumulate (e.g., Kidwell et al. 1977; Labrador et al. 1999; Kelleher et al. 2012). The strong accumulation of this Neptune element suggests that this element is particularly aggressive. Because naturally occurring sexually and asexually reproducing hybrid lineages have repeatedly evolved and can be compared with new lineages produced in the lab, we further propose that the *Fundulus* complex is an excellent system in which to disentangle hybridization-driven and asexuality-mediated effects on TE dynamics in the future.

## Data availability

Raw reads are available on SRA under PRJNA862733.

## Acknowledgements

We would like to thank Daniel Barbash, Mathieu Hénault, Karel Janko and France Dufresne for providing critical feedback. Part of the computations were enabled by resources in project NAISS 2024/22-720 provided by the National Academic Infrastructure for Supercomputing in Sweden (NAISS) at UPPMAX, funded by the Swedish Research Council through grant agreement no. 2022-06725.

