## Supplementary material for "Accumulation of a biparentally-inherited Neptune transposable element in natural Killifish hybrids (*Fundulus diaphanus* × *F. heteroclitus*)"

**Supplementary Tables**

**Table S1. Weight and measurements of classic landmarks for all individuals.** 13-17 is from the anterior insertion of the dorsal fin to the dorsal end of the caudal peduncle, 16-18 is from the posterior insertion of the anal fin to the ventral end of the caudal peduncle, and 17-18 is the peduncle height. All individuals were females.

| **ID** | **Weight (g)** | **13-17 (cm)** | **16-18 (cm)** | **17-18 (cm)** |
| --- | --- | --- | --- | --- |
| POR20-02 | 4.1 | 3.3 | 2 | 0.64 |
| POR20-03 | 3.8 | 2.35 | 1.7 | 0.7 |
| POR20-05 | 6.1 | 2.7 | 1.7 | 0.9 |
| POR20-06 | 2.85 | 2.2 | 1.5 | 0.72 |
| POR20-09 | 5.2 | 2.9 | 1.8 | 1 |
| POR20-10 | 6.16 | 3.15 | 1.8 | 0.8 |
| POR20-12 | 6.15 | 2.55 | 1.8 | 1 |
| POR20-13 | 2.5 | 2 | 1.45 | 0.65 |
| POR20-14 | 5.5 | 2.7 | 1.6 | 1 |
| POR20-15 | 7.2 | 2.8 | 1.9 | 1 |
| POR20-19 | 4.56 | 3.2 | 1.9 | 0.8 |
| POR20-20 | 3.69 | 3.1 | 1.6 | 0.75 |
| POR20-21 | 3.16 | 2.1 | 1.4 | 0.9 |
| POR20-22 | 3.54 | 2.05 | 1.4 | 0.8 |

**Table S2. Multi-locus genotypes for all individuals**. Fhe113 does not amplify the *F. diaphanus* allele, so hybrid individuals appear homozygous.

| **ID** | **mtDNA** | **FhCA-1** | | **FhCA-21** | | **Fhe57** | | **FhATGB103** | | **Fhe113** | | | **Species** |
| --- | --- | --- | --- | --- | --- | --- | --- | --- | --- | --- | --- | --- | --- |
| POR20-02 | *F. diaphanus* | 174 | 174 | 151 | 151 | 179 | 179 | 332 | 344 | - | - | *F. diaphanus* | |
| POR20-03 | *F. diaphanus* | 142 | 180 | 151 | 175 | 144 | 182 | 345 | 350 | 209 | 209 | Hybrid | |
| POR20-05 | *F. diaphanus* | 142 | 180 | 151 | 175 | 144 | 182 | 344 | 345 | 209 | 209 | Hybrid | |
| POR20-06 | *F. diaphanus* | 142 | 180 | 151 | 175 | 144 | 182 | 345 | 350 | 209 | 209 | Hybrid | |
| POR20-09 | *F. diaphanus* | 142 | 180 | 151 | 175 | 144 | 182 | 345 | 350 | 209 | 209 | Hybrid | |
| POR20-10 | *F. diaphanus* | 174 | 174 | 151 | 151 | 179 | 179 | 335 | 344 | - | - | *F. diaphanus* | |
| POR20-12 | *F. heteroclitus* | 144 | 144 | 175 | 175 | 144 | 144 | 375 | 394 | 216 | 216 | *F. heteroclitus* | |
| POR20-13 | *F. diaphanus* | 142 | 180 | 151 | 175 | 144 | 182 | 345 | 350 | 209 | 209 | Hybrid | |
| POR20-14 | *F. diaphanus* | 142 | 180 | 151 | 175 | 144 | 182 | 345 | 350 | 209 | 209 | Hybrid | |
| POR20-15 | *F. diaphanus* | 142 | 180 | 151 | 175 | 144 | 182 | 345 | 350 | 209 | 209 | Hybrid | |
| POR20-19 | *F. diaphanus* | 174 | 174 | 151 | 151 | 179 | 179 | 332 | 344 | - | - | *F. diaphanus* | |
| POR20-20 | *F. diaphanus* | 174 | 174 | 151 | 151 | 179 | 179 | 347 | 350 | - | - | *F. diaphanus* | |
| POR20-21 | *F. heteroclitus* | 144 | 144 | 175 | 175 | 144 | 149 | 375 | 400 | 209 | 212 | *F. heteroclitus* | |
| POR20-22 | *F. heteroclitus* | 142 | 142 | 175 | 175 | 144 | 144 | 396 | 418 | 209 | 214 | *F. heteroclitus* | |

**Table S3. Sequencing summary.** Genome coverage was estimated by dividing the total read length (number of reads X 150bp) by the estimated genome size (Dawley *et al.* 1992). *F. heteroclitus* has an estimated genome size of 1.33 Gb whereas *F. diaphanus* has an estimate of 1.48 bp. The hybrid genome size was considered to be intermediate at 1.40 Gb.

| **ID** | **Species** | **number of reads** | **Genome Coverage** |
| --- | --- | --- | --- |
| POR20-02 | *F. diaphanus* | 96,957,478 | 9.8 |
| POR20-03 | Hybrid | 113,635,650 | 12.1 |
| POR20-05 | Hybrid | 114,423,954 | 12.2 |
| POR20-06 | Hybrid | 118,412,098 | 12.7 |
| POR20-09 | Hybrid | 91,534,340 | 9.8 |
| POR20-10 | *F. diaphanus* | 98,612,536 | 10 |
| POR20-12 | *F. heteroclitus* | 73,223,826 | 8.3 |
| POR20-13 | Hybrid | 87,983,898 | 9.4 |
| POR20-14 | Hybrid | 94,260,146 | 10.1 |
| POR20-15 | Hybrid | 116,836,360 | 12.5 |
| POR20-19 | *F. diaphanus* | 87,469,854 | 8.9 |
| POR20-20 | *F. diaphanus* | 84,252,256 | 8.6 |
| POR20-21 | *F. heteroclitus* | 87,373,042 | 9.9 |
| POR20-22 | *F. heteroclitus* | 107,407,738 | 12.1 |

**Table S4. Repeat content.** Genome proportion was calculated by summing up the abundance value of each divergence bin per repeat category (as quantified by RepeatMasker) and dividing that value by the estimated genome size for each species (Dawley et al. 1992). *F. heteroclitus* has an estimated genome size of 1,330,080,000 base pairs (bp) whilst *F. diaphanus* is at an estimate of 1,476,780,000 bp. The hybrid genome was considered to be halfway between the parental species and was thus estimated at 1,403,430,000 bp. Annotation based on Figure S1.

|  |  | Genome proportion (%)  (bp) | | | | |
| --- | --- | --- | --- | --- | --- | --- |
| Category | # clusters | *F. diaphanus* | *F. heteroclitus* | Hybrids  observed (o) | Hybrids  expected (e) | Hybrids  Difference (o-e) |
| rDNA | 2 | 0.31  (4,554,603) | 0.48  (6,358,479) | 0.19  (2,760,076) | 0.39  (5,456,541) | -0.2  (2,696,465) |
| Unknown | 711 | 15.44 (227,966,393) | 14.71  (195,701,327) | 15.32  (215,063,088) | 15.09  (211,833,860) | 0.23  (3,229,228) |
| DIRS  (non-LTR) | 8 | 0.06  (852,950) | 0.09  (1,156,531) | 0.09  (1,233,775) | 0.07  (1,004,741) | 0.02  (229034) |
| LINE  (non-LTR) | 184 | 8.36  (123,445,704) | 7.28  (96,891,723) | 8.06  (113,115,172) | 7.85  (110,168,714) | 0.21  (2,946,458) |
| Other LTR | 19 | 0.47  (6,966,120) | 0.36  (4,787,409) | 0.42  (5,949,208) | 0.42  (5,876,765) | 0  (72,443) |
| Maverick (DNA) | 16 | 0.11  (1,638,762) | 0.07  (934,789) | 0.10  (1,424,009) | 0.09  (1,286,776) | 0.01  (137,233) |
| Penelope  (non-LTR) | 16 | 0.33  (4,865,093) | 0.37  (4,968,926) | 0.36  (4,989,836) | 0.35  (4,917,010) | 0.01  (72,826) |
| Ty3 (LTR) | 1 | 0.59  (8,749,211) | 0.70  (9,323,783) | 0.73  (10,208,067) | 0.64  (9,036,497) | 0.09  (1,171,570) |
| Retrovirus (ERV) | 4 | 0.18  (2,607,938) | 0.04  (555,913) | 0.11  (1,593,618) | 0.11  (1,581,926) | 0  (11,692) |
| Unclassified repeats | 84 | 3.05  (45,088,788) | 2.69  (35,754,185) | 3.45  (48,376,237) | 2.88  (40,421,487) | 0.57  (7,954,750) |
| Satellite | 33 | 1.49  (22,045,953) | 2.51  (33,437,030) | 2.20  (30,847,151) | 1.98  (27,741,492) | 0.22  (3,105,659) |
| Total | 1078 | 30.39  (448,781,514) | 29.30  (389,870,096) | 31.04  (435,560,237) | 29.88  (419,325,805) | 1.16  (16,234,432) |

**Supplementary figures**

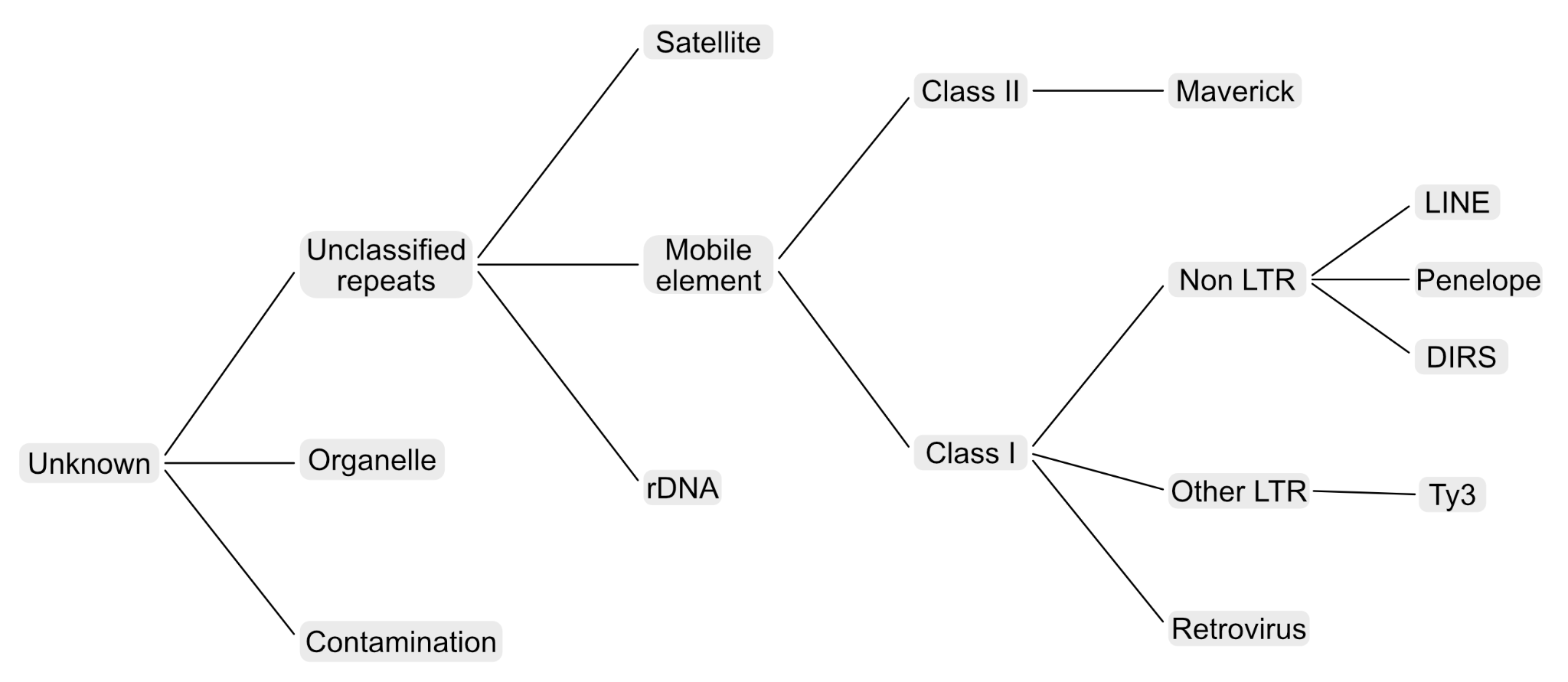

**Figure S1.** **Annotation decision tree.** Simplified and adapted from Novák et al. 2020.

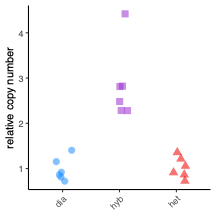

**Figure S2. Copy number of the reverse transcriptase encoded by repeat.CL27 based on qPCR.** The copy number of the reverse transcriptase is relative to the citrate synthase gene.

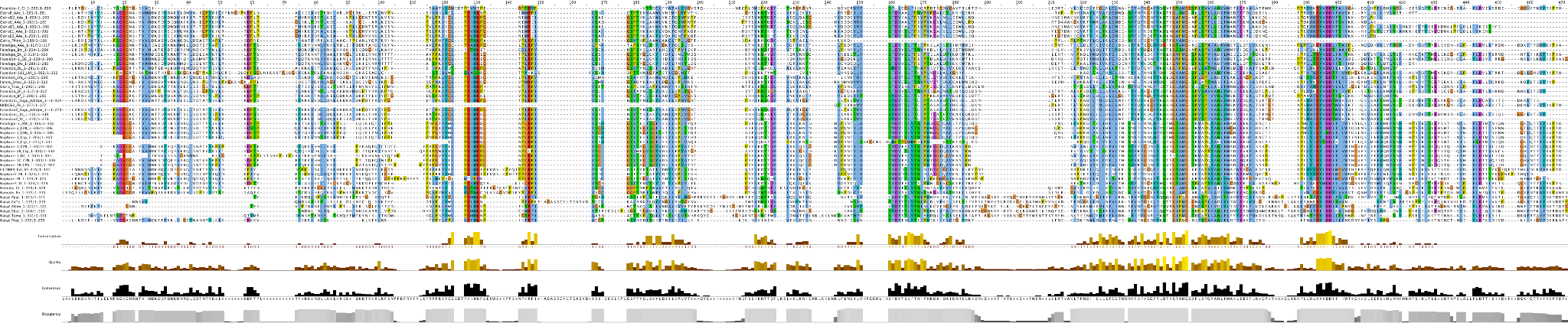

**Figure S3.** Multiple sequence alignment of *Penelope*-like elements and ORF9 from repeat.CL27.

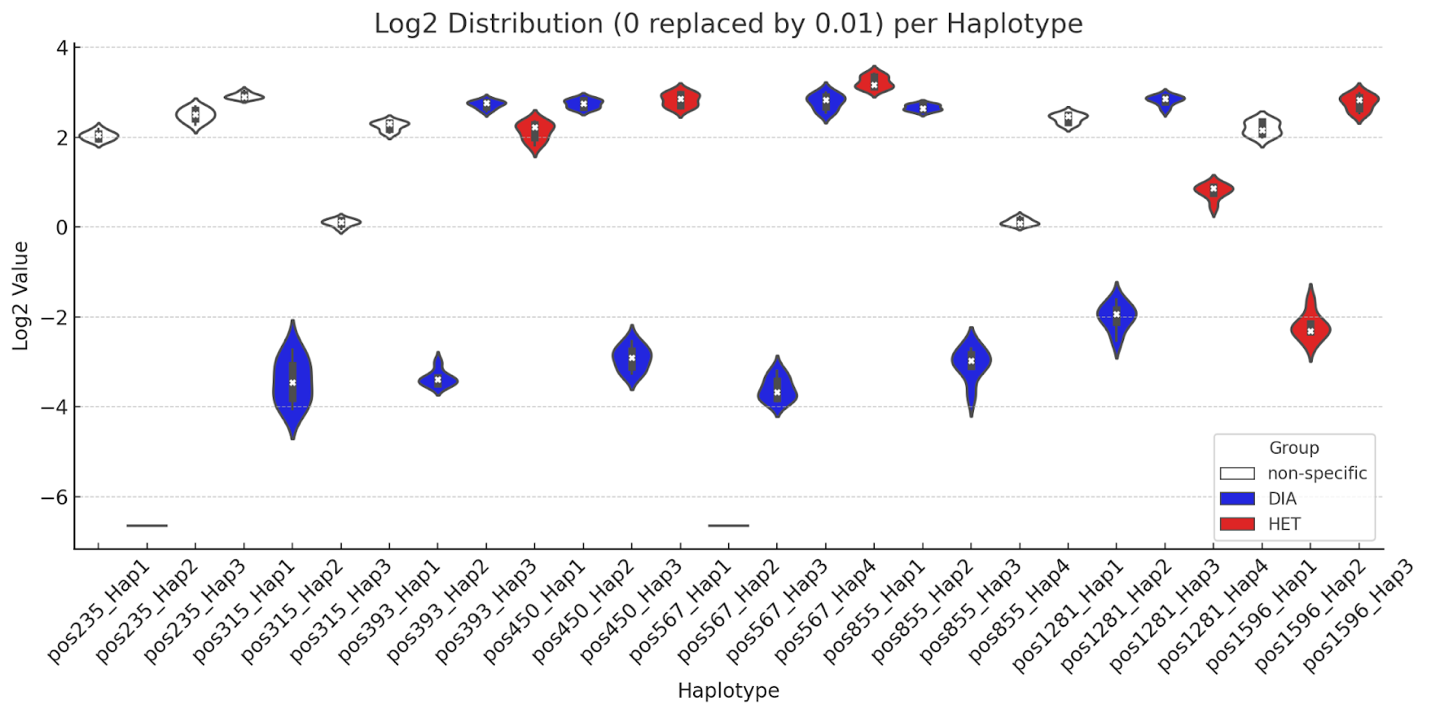

**Figure S4. Deviation from the expected frequency of haplotypes in the CL27.repeat with 15 or more nucleotides.** Observed frequency for each hybrid individual was divided by the expected frequency calculated as the average of both parental species. Values were transformed as log2, showing positive values when observed frequencies are higher than expected and negative values when observed frequencies are lower than expected. Colors of the violin plots indicate the parental species where each haplotype is specific: blue for *F. diaphanus* and red for *F. heteroclitus*. When there were no observations, we replaced 0 by 0.01.
